# Defending as a unit: sex- and context-specific territorial defence in a duetting bird

**DOI:** 10.1101/2020.07.29.226167

**Authors:** Lucia Mentesana, Maria Moiron, Ernesto Guedes, Enzo Cavalli, Bettina Tassino, Nicolas M. Adreani

## Abstract

Behaviours such as territorial defence represent functionally integrated traits that underlie multiple behavioural variables such as physical and acoustic responses. Characterizing the multivariate structure of such traits is fundamental to understand their evolution. In bird species that form stable pair bonds and are territorial year-round, both sexes are expected to defend their territory; however, the role that each sex plays in defending their shared territory remains largely unknown. Evidence for the sex-roles during territorial defence is mixed and sex- and context-specific characterizations of territorial defence embracing the multivariate nature of the trait are currently lacking. Here we investigated sex- and context-specific variation in a hypothesised latent variable called “territorial defence” and tested whether duets were part of territorial defence in a wild population of rufous hornero (*Furnarius rufus*). To do so, we combined a simulated territorial intrusion approach during nest building and provisioning contexts with a structural equation modelling approach. Our results showed that, in males and females, the six measured behavioural variables were linked by a single latent trait, territorial defence, in both contexts. Flights over the decoy and duet songs were equally good proxies of territorial defence. Although males were defending more the territory than females, pair members showed a positive correlation in their behaviour. The structural equation modelling framework enabled us to capture a complex correlation pattern among behavioural variables, expanding upon a classic body of research on territorial defence. Thus, the combination of classical behavioural approaches with sophisticated statistical analyses brings new exciting possibilities to the field of behavioural ecology.

**Significance statement:** Territorial defence is a key behaviour in territorial species as it plays a major role in an individual’s reproductive success and survival. Additionally, territorial defence has been proposed as one possible evolutionary driver of duetting behaviour, one of the most fascinating vocal behaviours in birds. As behaviours are evolutionary characters, they must be studied in a multivariate framework. In this study we focused on characterizing territorial defence during a simulated territorial intrusion in an integrative manner using a classical territorial intrusion framework. We did so in male and female rufous horneros (Aves: *Furnaridae*) across two breeding contexts, while simultaneously testing theoretical predictions about the role of duetting behaviour as key part of territorial defence. Overall, our study provides for the first time a sex- and context-comparison of the multivariate, latent variable “territorial defence” in duetting birds, while highlighting the potential of combining field behavioural approaches with structural equation modelling.

## Introduction

Behaviours, such as territorial defence, are typically studied by simultaneously measuring different observable variables (e.g. “number of attacks”, “latency of response”, “proximity” measures, and/or “vocal responses”; Wingfield 1994; Bollen 2002). Extensive empirical research over the last decades has focused on analyzing male and, to a lesser extent, female territorial defence behaviour using different approaches. Evidence generally shows that multiple components of an animal’ defence response in a territory intrusion are intercorrelated (e.g. Huntingford 1976; Sprenger et al. 2012). When these multiple behaviours are functionally related, they might be considered expressions of a single evolutionary character (Araya-Ajoy and Dingemanse 2013). Their study should therefore not be addressed by means of bivariate correlations but integrate the multivariate nature of the behaviour by quantifying unobserved, biologically-relevant latent variables (Houle et al. 2011; Carter and Feeney 2012; Araya-Ajoy and Dingemanse 2013). One way to characterize “territorial defence” as an evolutionary character while also quantifying which of the observed behavioural variables should be considered expressions of it, is to apply a Structural Equation Modelling approach (SEM; e.g. Card and Little 2007). This statistical framework allows to explore complex correlation patterns among multiple behavioural variables, and to test *a priori* defined hypotheses of how multiple observed behavioural variables are linked by the unmeasured latent variable (Araya-Ajoy and Dingemanse 2013).

Territorial defence behaviour has been widely studied in diverse organisms from insects to several groups of vertebrates (reviewed in Smith and Blumstein 2008), likely because of its impact on fitness (Stamps and Krishnan 1997; Smith and Blumstein 2008). During territory defence, aggressive interactions can be beneficial for both males and females because an intrusion of a conspecific into the breeding territory might, for example, lead to loss of limited resources (Stamps and Krishnan 1997; Garcia and Arroyo 2002). Nevertheless, the sex-specific contribution to territory defence differs among species according to variation in mating systems and parental care (Emlen and Oring 1977; Clutton-Brock and Vincent 1991; Owens and Thompson 1994). In birds, among those species that form stable pair bonds and are territorial year round, it is expected that males and females equally contribute to territory defence (Greenberg and Gradwohl 1983). In line with this prediction, in dot-winged antwrens (*Microrhopia quixensis*) and in the purple-crowned and red-backed fairy-wrens (*Malurus coronatus, Malurus melanocephalus*) both sexes contribute to the same extent to defend their territory (Greenberg and Gradwohl 1983; Hall and Peters 2008; Dowling and Webster 2016). However, these findings were not observed in other antbird species (*Phaenostictus mcleannani*, Willis 1972; *Cercomacra tyrannina*; Morton and Derrickson 1996; *Hylophylax naevioides*; Bard et al. 2002; *Myrmeciza longipes*; Fedy and Stutchbury 2005), in the zenaida dove (*Zenaida aurit*a; Quinard and Cézilly 2012) and in the rufous hornero (*Furnarius rufus;* Diniz et al. 2018), where males engaged more in defensive interactions than females. In addition, whilst intensity of territorial defence can be positively correlated within pairs in some species (especially on those that duet e.g. Logue 2005; Hall and Peters 2008), in others the opposite relationship is true (e.g. zenaida dove; Quinard and Cézilly 2012). Therefore, the generality of sex-specific territorial defence as well as its intensity from both members of a pair remains poorly understood in species that are socially monogamous and territorial year-round. Furthermore, it also remains largely unknown whether the same observed behavioural variables characterize the latent variable territorial defence in males and females and across different breeding contexts. For instance, it is still an open question whether territorial defence is equally characterized by physical and vocal behaviours in both sexes and across contexts. Studies describing territorial defence as a latent variable using SEM and explaining broader patterns of territorial defence across sexes or contexts, will help to shed light on these questions and further our understanding of the evolution of male and female defence of territory.

While physical displays are considered the main defensive responses, the role of vocal displays as defensive signals remains under discussion (see Searcy and Beecher 2009; Naguib and Mennill 2010). Among the vocal displays that take place during agonistic interactions, perhaps the most fascinating one is duetting – occurring in around 18% of avian species worldwide (Tobias et al. 2016). Duets are defined as coordinated vocal interactions between two individuals - usually a male and female of a pair - that occur with a given temporal precision (Farabaugh 1982; Hall 2004). Duets are hypothesised to represent an important component of territorial defence (Langmore 1998; Hall 2004). In particular, the “joint territorial defence” hypothesis, proposed as one evolutionary driver of duetting behaviour (Wickler and Seibt 1980), postulates that duets allow pairs to cooperatively defend resources from conspecific intruders (Robinson 1949; reviewed by Hall 2004). A central prediction of the “joint territorial defence” hypothesis is that duets are threatening signals, stronger than solo songs (Hall 2004). To date, few studies investigated duetting in the context of territorial defence across different life-history stages (Topp and Mennill 2008; Odom et al. 2017; Quirós-Guerrero et al. 2017; Sosa-López et al. 2017; Diniz et al. 2018). The few that did so used mainly three methods: i) context criterion (i.e. which compares responses towards acoustic stimuli that represent different contexts, like only male/female solo songs, only duet songs or only heterospecific songs; e.g. Dowling and Webster 2016), ii) response criterion (i.e. which compares responses with and without a playback stimulus e.g. Hall and Peters 2008), and iii) correlation methods (i.e. which applies correlation techniques and principal component transformations on behavioural data, e.g. Kolof and Mennil 2013). However, none of the abovementioned methods allow to test the role that duets play in the context of territorial defence while embracing the multivariate nature of this traits as a latent variable.

We conducted simulated territorial intrusions (STI) in the territory of focal pairs of rufous horneros during two contexts in the breeding season: nest building and chick provisioning. The rufous hornero, hereafter hornero, is a single brooded furnarid bird species that is widely distributed throughout southern South America (Fraga 1980). Horneros are territorial year round, socially monogamous (Fraga 1980; Diniz et al. 2018) and both members of the pair are involved in defending their territory (Fraga 1980; Diniz et al. 2018). Indeed, all breeding behaviours studied in horneros so far are performed in an equitable and coordinated manner between sexes, such as incubation, parental care-related activities and even territorial defence in non-breeding context (Fraga 1980; Massoni et al. 2012; Diniz et al. 2020). Also, previous studies on this species suggest that duets have a territorial function (Diniz et al. 2018, 2019, 2020). However, these studies were either acoustic-centered (Diniz et al. 2018, 2019) or carried out during a non-breeding season (Diniz et al. 2020), and none of them considered the multivariate nature of territorial defence.

The main goals of our study were first to characterize the multivariate nature of territorial defence in male and female horneros, and to test whether duets were indeed part of the defence displays during a territorial intrusion. Our second goal was to quantify phenotypic variation across sexes, breeding contexts and pair members. Our third goal was to evaluate the level of coordination between sexes during territorial defence. To do so, we constructed a series of structural equation models where we tested three hypotheses of potential associations among the behavioural variables: model 1 hypothesised that each territorial defence behaviour is independent and not part of a functional unit or evolutionary character; model 2 hypothesises that one latent variable, “territorial defence”, underlies the relationships between all behavioural variables; and model 3 hypothesises that all behavioural variables except number of duet songs are linked by the latent variable “territorial defence”. These models were therefore specifically constructed to test the “joint territorial defence” hypothesis (Hall 2009). According to this hypothesis, for our first aim, we predicted duets to be part of the latent trait ‘territorial defence’ and to be more relevant than solo songs. We also predicted that territorial defence will be characterized by the same behavioural traits (i.e. number of duets, number of flights over the decoy, times spent within 5m of the decoy, number of solo songs, time spent on nest) in males and females. Second, we predicted that in our STIs males would defend more their territories than females. This was based on the notion that, although in neotropical birds there is mixed evidence for the sexual difference in territorial defence, a recent study reported that male horneros engaged more in defending their territories than females (Diniz et al. 2018). Further, because in horneros territory take-over is expected to be a stronger driver of aggression than paternity loss (i.e. extra-pair paternity levels are ∼ 3%; Diniz et al. 2019), we predicted higher levels of territory defence earlier in the nest building than in the provisioning context (see also Demko and Mennill 2018). Finally, we predicted both members of the pair to positively correlate their territory defence behaviors (e.g. Diniz et al. 2020).

## Materials and Methods

### Field site and experimental procedures

We studied pairs of horneros in two periods during 2016 on the campus of INIA “Las Brujas” (National Institute of Agricultural Research), department of Canelones, Uruguay (34°40’ S, 56°20’ W; 0-35 m a.s.l.). Behavioural assays were carried out during “nest-building” (i.e. when pairs were observed finishing their nests and females were in their fertile period; August 23^rd^ – September 27^th^), and “provisioning” periods (i.e. when pairs were observed feeding their young; November 7^th^ – December 6^th^). Overall, we observed 39 males and 38 females during nest building and 25 males and 24 females during provisioning. Each pair was tested only one time (i.e. either during the nest-building or during the provisioning period). It was not possible to record data blind because our study involved focal animals in the field. All the behavioural assays were performed between 07h00 and 13h00. We quantified territorial defence behaviours by performing simulated territorial intrusions (STI) in the territory of a focal pair. Once the focal pair was identified, a stuffed decoy of a hornero together with playback sounds was presented 10 metres away from the pair’s nest. Using two decoys could have been more realistic; however, we could not do this for ethical reasons. Every STI playback lasted 20 minutes and consisted of randomly selected stimuli from a pool of ten male solo songs, ten duets and ten audio files containing 7-15 seconds of silence. The auditory stimuli for each STI were randomly selected to avoid pseudo-replication of the acoustic component across territories (e.g. Apfelbeck et al. 2011), to avoid a behavioural bias towards specific songs or duets, and to elicit comparable behavioural responses across territories. Our approach hinged on the notion that horneros are suboscines and do not learn their vocalizations (Freeman et al. 2017). This is, compared to oscines, the acoustic variability of songs and duets across individuals is low (Freeman et al. 2017). We played silence tracks of different length to avoid habituation in the focal birds. All playback sounds were “wav” files recorded in Uruguay and were obtained from the database of Xeno-canto (www.xeno-canto.org) and normalized in amplitude. All the sounds were broadcasted from a single speaker (Douglas and Mennil 2010). Although a multiple-speaker approach would have been more realistic, because horneros frequently perform their duets while close to each other, it is unlikely that our set-up introduced a bias in the STIs. We simultaneously recorded the behaviour of each individual of the pair during the 20 minutes of STI (mean ± SE; 19.96 ± 0.23 min). Two observers performed the observations from a distance of 15 metres using digital voice recorders (Philips VoiceTraicer DVT1200 and Olympus Digital Recorder VN-733 PC). The focal bird was randomly assigned to each observer. The following measures were recorded: 1) response latency (time between start of playback and first approach within 10 metres from the dummy), 2) time spent within five metres of the decoy, 3) average time spent on the nest during a visit, 4) number of solo songs, 5) number of duet songs, and 6) number of flights over the decoy (i.e. flights directed to and over the decoy). Regarding the variable ‘number of duets’, we initially considered the fact that both males and females can initiate the duet (Diniz et al, 2018). However, during our STIs there were seldom cases in which the female initiated the duet, and none were during nest building. For this reason, we only considered the number of duets as a joint variable across sexes in our models. Additionally, the solo songs in males represent instances in which females decided not to join in the duet. The sex of each bird could be determined from the acoustic signature of each individual because the vocal contribution of each sex in the duet is dimorphic (Roper 2005). As part of a different project, birds were captured after the STI and the sex was verified by PCR (sex was correctly assigned by the observers in 96.3% of the cases for those individuals to whom the sex could be assigned acoustically and were trapped in the nets; for details see Adreani et al. 2018).

### Statistical analyses

#### (i) Structural equation modelling

First, we applied a structural equation modelling (SEM) approach to study three *a priori* hypotheses of relationships among the six behavioural variables quantified during the simulated territorial intrusion (i.e. response latency, time spent within five metres of the decoy, time spent on the nest, number of solo songs, number of duet songs, and number of flights over the decoy; Fig. 1). Of the three models for each sex and context, model 1 represents a (biologically unrealistic) “null” expectation (i.e. each defensive behaviour is independent and not part of a functional unit or evolutionary character); model 2 hypothesises that one latent variable, “territorial defence”, underlies the relationships between the six behavioural variables; and model 3 hypothesises that all behavioural variables except number of duet songs are linked by the latent variable “territorial defence”. While more complex structural models could be constructed (i.e. including trade-offs between behavioural variables), the present framework is the one that allows for a straightforward testing the “joint territorial defence” hypothesis (Hall 2009). We estimated each structural equation model separately for males and females because we only have one measure of the number of duet songs from a single pair and, therefore, it is not possible to disentangle the sex-differences in number of duets on the latent variable territorial defence. We also estimated each model separately for the two breeding contexts (i.e. nest building *vs.* provision context). The formulation of these four different sets of models allowed us to qualitatively assess whether there were differences between sexes and breeding contexts (nest building and provisioning) in the structure and strength of the hypothesized latent variable. Therefore, besides characterizing the latent variable structure, we were also interested in qualitatively investigating differences between sexes and contexts in path loadings across models (i.e. whether behavioural traits maintain their rank differences among path loadings). We also constructed a single model for each sex in both breeding contexts, where the 12 different behavioural variables were modelled simultaneously. However, we decided to present here the separated models, one for each breeding context, because the full model (i.e., with the 12 variables) is likely over-parametrized (i.e., there was a compromise between the complexity of the SEM models fitted and the number of observations given the number of variables tested in each SEM). See Supplementary Material for further details on the full model (Table S1-2 and Fig S1).

**Fig. 1.**
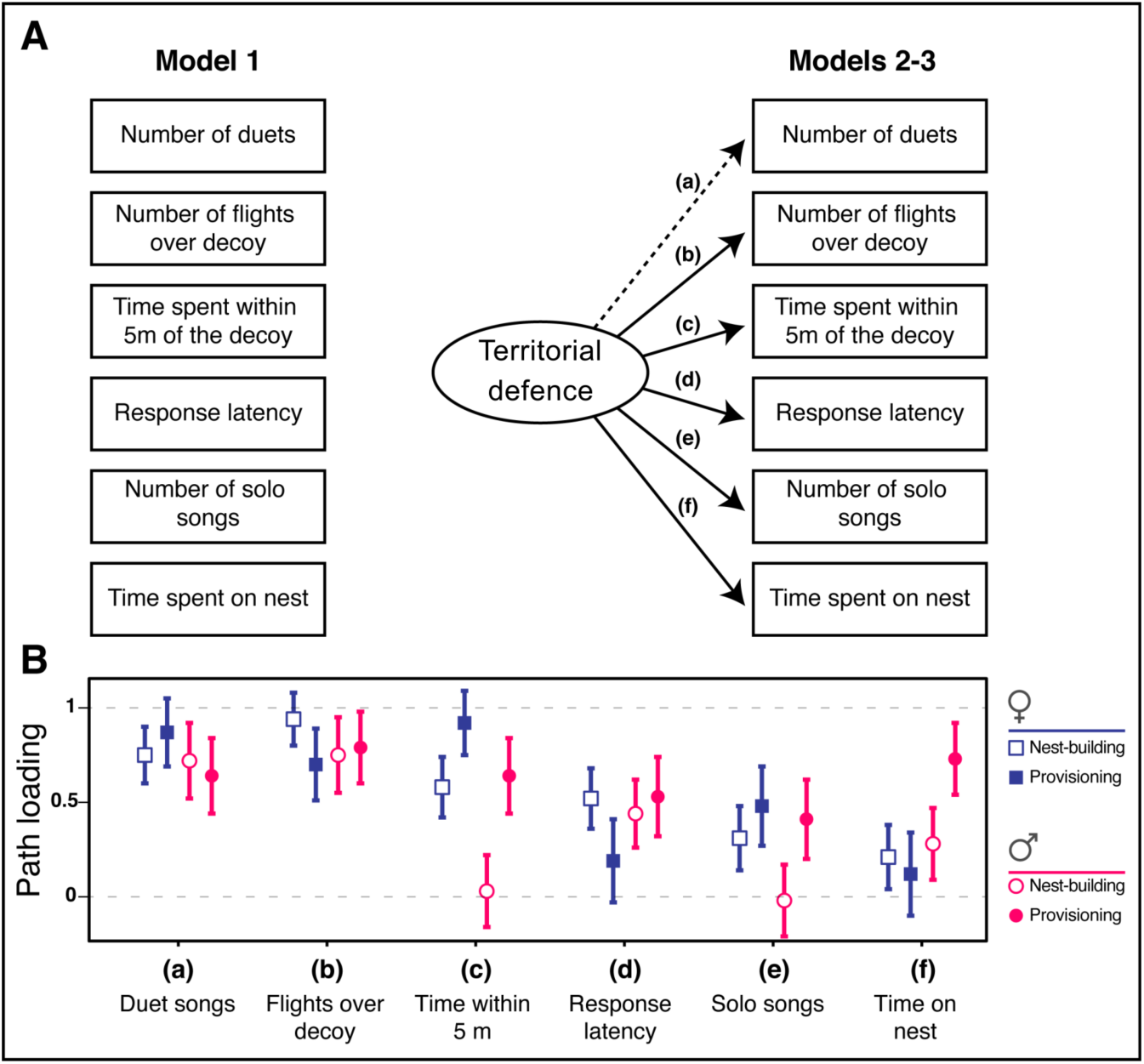
(A) Three models (hypotheses) explaining the correlation structure among behavioural variables assayed during a simulated territory intrusion in the nest building and provisioning context in wild horneros. Model 1 hypothesises trait independence (null model); model 2 hypothesises a latent variable (“territorial defence”) linking all behavioural variables, whereas model 3 hypothesises a latent variable (“territorial defence”) linking all behavioural variables except “number of duets”. Unidirectional arrows represent directional relationships between traits. Solid lines represent relationships present across all models; and the dashed line represents a relationship expressed in a specific model structure. Path “a” is only active in model 2. (B) Path loadings of model 2 for males and females in both breeding contexts. Squares and circles represent the estimated mean, and error bars represent the standard error of the mean

To test the relative fit of each alternative biological hypotheses, we first estimated the matrix of phenotypic correlations of all the behavioural variables for each combination of sex and breeding context. The correlation matrix was constructed using Spearman coefficients obtained with the R package “stats” in R v. 3.3.3 (R Core Team 2013). Data on “response latency” were missing for one out of 38 individual females during nest building and data for “time spent on the nest” were missing for two out of 24 females during provisioning. We assigned the average population phenotypic value of each trait to those individuals with missing values (note that a “complete-case analysis” did not change our findings, results not shown). We then implemented each among-trait correlation matrix in the R-package “sem” and tested the different SEM hypotheses. We statistically compared each model’s fit using the Akaike Information Criterion for small sample sizes (AICc; Burnham and Anderson 2004) and evaluated their relative support based on AICc differences relative to the best-fitting model (ΔAICc). We also present values for the Goodness of Fit Index (GFI), which represents the proportion in the improvement of the overall fit of a given hypothesised model compared to the independence model. GFI values range from 0 (poor fit) to 1 (perfect fit), being considered as satisfactory when it is higher than 0.9.

#### (ii) Univariate mixed-effect models

As a second step, we investigated sources of variation in territorial defence behaviour in our population of horneros using a series of univariate mixed models. This step was necessary because we were also interested in investigating sex-differences in territorial defence. However, as we only had a single measure of number of duets per pair, we could not test for sources of variation in territorial defence using our multivariate SEM approach. Thus, we ran two univariate mixed-effect models fitting number of flights over the decoy and number of duets as the response variables, respectively. Though we had measured various potential proxies of territorial defence (detailed above), we used number of flights over decoy and number of duet songs because they consistently had the highest value in path loading across all models (for a further discussion on the rationale of this approach, see Araya-Ajoy and Dingemanse 2013). Breeding context (nest building vs. provisioning), sex (male vs. female) and their interaction, time of the day (i.e. moment of the day when the territorial intrusion was simulated, expressed in decimal fractions of hours after sunrise and mean centred), and observer identity (observer 1 vs. 2) were included as fixed effects in the univariate mixed-effect models. Time of day was mean centred, such that the fixed-effect intercept of the model was estimated for the behavioural trait on the average time (following Dingemanse and Dochtermann 2013). In the model with “number of flights over the decoy” as a response variable, we fitted random intercepts for pair identity (“Pair identity”; *n* = 63 levels). In the model with “number of duet songs”, we did not include this random effect because we did not have repeated measures of duet song frequency for the same pair identity. Both response variables, number of flights over the decoy and number of duet songs, were modelled with Poisson errors with a log-link function. In both models, we included an observation level random effect to account for over-dispersion (Harrison 2014). The analyses were performed using the R packages “lme4” (Bates et al. 2014) and “arm” (Gelman and Yu-Sung 2015). We used the “sim” R function to simulate posterior distributions of the model parameters. Based on 5000 simulations, we extracted the mean value and 95% Credible Intervals (CrI) of the posterior distributions. Model fit was assessed by visual inspection of the residuals. Assessment of statistical support was obtained from the posterior distribution of each parameter (Zuur 2016). We considered an effect “strongly supported” if zero was not included within the 95% CI, and “moderately supported” if the point estimate was skewed away from zero while its 95% CI simultaneously overlapped zero. Estimates centred on zero were viewed as strong support for the absence of an effect.

#### (iii) Correlation between male and female contribution to territorial defence

We studied the correlation pattern between male and female territorial defence to investigate whether defence response within a pair was positively correlated. To do so, we estimated the Spearman correlation coefficient of “number of flights over the decoy” between males and females from a single pair. This correlation was calculated by including in our analysis all pairs observed during both contexts (*n* = 91 pairs), given that we found no differences in the number of flights over the decoy between the nest building and provisioning contexts (see below). The correlation test was performed using the R-package “stats” in R v. 3.3.3 (R Core Team 2013).

## Results

### (i) Territorial defence as a latent trait and the role of duetting

The behavioural variables assayed during the simulated territorial intrusion (i.e. response latency, time spent within five metres of the decoy, time spent on the nest, number of solo songs, number of duet songs, and number of flights over the decoy) were, to a varying extent, correlated with each other; both across sexes and contexts (Table S3, S4). Overall, horneros with shorter latency of response to the territorial intrusion spent more time within five metres of the decoy and on the nest, sang more solo and duet songs, and flew more often over the decoy, suggesting the existence of the hypothesised latent variable “territorial defence” linking the six behavioural variables.

AICc model comparison identified the SEM model 2 as the best one (among the models we fitted) explaining the structure of the phenotypic variables across the four different set of models (Table 1). Model 2 represented an overarching latent variable (“territorial defence”) linking the expression of all behavioural variables, including the number of duet songs (Fig. 1A). Furthermore, the number of duet songs together with the number of flights over the decoy had consistently the highest values of path loading in males and females for both breeding context (Fig. 1B, Table S5). We thus considered them both equally good predictors for territory defence in horneros. Furthermore, a standard index of model fit (“Goodness of Fit Index”) considered satisfactory our best fitting model across all models (i.e., GFI values for Model 2 were around or above 0.90 across all models, Table 1).

**Table 1.**
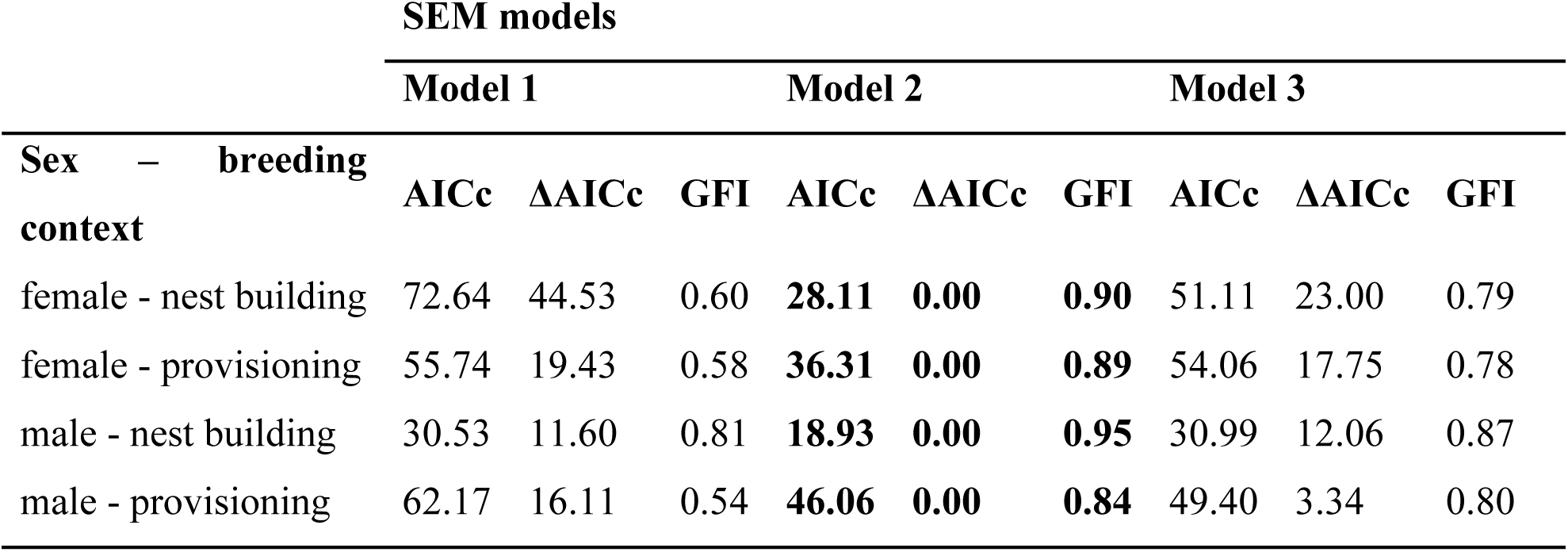
Results of model comparison using Akaike Information Criterion for small sample sizes (AICc) values to compare our three candidate models. Smaller AICc values are given to models that better fit the data. Models whose AICc values differ from that of the top model (ΔAICc) by more than 2 are considered to lack explanatory power relative to the top model. We also present values for the Goodness of Fit Index (GFI). The best-supported hypothesis is printed in boldface

### (ii) Effect of sex, breeding context and time of the day on territorial defence

We did not find strong evidence that horneros differed in the number of duet songs between the nest building and provisioning context. While the effect size is moderately supported, the evidence is weak due to large uncertainty (Table 2; Fig. 2A). We did not find differences in the number of duets explained by time of the day (Table 2). Regarding the number of flights over the decoy, males were on average defending more (i.e. flew more times over the decoy) than females during both breeding contexts (Table 2; Fig. 2B). However, we again found weak evidence that horneros differed in the number of flights over the decoy between the nest building and provisioning context, and there were also no sex-specific differences between the two breeding contexts (i.e. the effect sizes are relatively large but estimated with large uncertainty, therefore the support is moderate; Table 2, Fig. 2B). Furthermore, we observed moderate effects of time of day and observer identity in our model (the estimates include zero in their 95% CrI, but the effect sizes are considerable).

**Table 2.** Sources of variation in “number of flights over the decoy” and “number of duets” in horneros. Breeding context (nest building vs provisioning), sex (female vs male) and their interaction; time of the day (hours after sunrise, mean centred); and observer identity (Observer 1 vs 2) were fitted as fixed effects. Pair identity and an observation-level parameter were fitted as random effects. Both response variables were modelled with Poisson error. We present estimates of fixed (β) and random (σ^2^) parameters with their 95% Credible Intervals (CrI) in brackets. The reference category for the categorical variable sex is “female”; for breeding context, “nest building”; and for observer identity is “observer 1”

| Fixed effects | Flights over decoy | Duets |
| --- | --- | --- |
| | $\beta$ (95% CrI) | $\beta$ (95% CrI) |
| Intercept | 2.33 (1.87, 2.78) | 2.44 (2.21, 2.69) |
| Sex | 0.58 (0.32, 0.83) | -- |
| Breeding context | -0.28 (-0.88, 0.31) | -0.11 (-0.52, 0.28) |
| Sex × Breeding context | 0.22 (-0.19, 0.64) | -- |
| Time of day | -0.19 (-0.45, 0.05) | -0.07 (-0.26, 0.13) |
| Observer Identity | -0.16 (-0.36, 0.04) | -- |
| Random effects | $\sigma^2$ (95%CrI) | $\sigma^2$ (95%CrI) |
| Pair Identity | 1.02 (0.75, 1.36) | -- |
| Observation-level parameter | 0.18 (0.14, 0.24) | 0.44 (0.33, 0.60) |

**Fig. 2.**
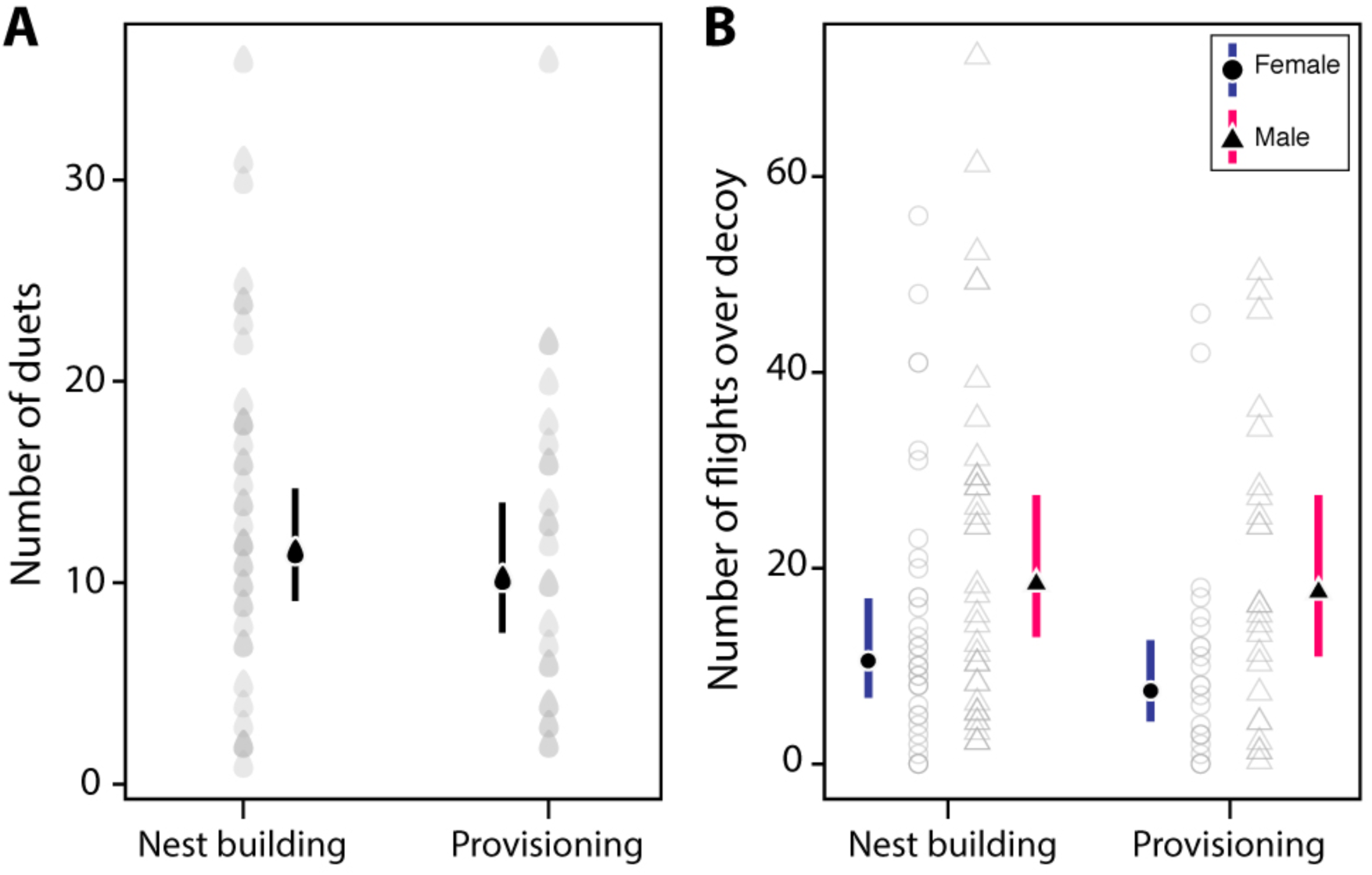
Number of (A) duets and (B) flights over decoy of hornero pairs during nest building and provisioning context. Grey symbols represent raw data. The mean estimates of the posterior distributions (black symbols) as well as the 95% credible intervals (error bars) are also shown

### (iii) Defending as a unit: correlation between male and female territorial defence

We investigated whether the defensive response to a territory intrusion of an individual was correlated with the response expressed by its partner. We found that territorial defence of males and females within pairs was strongly positively correlated (ρ = 0.67, p < 0.0001). Thereby, within a single pair, male and female had matching levels of defensive response (Fig. 3).

**Fig. 3.**
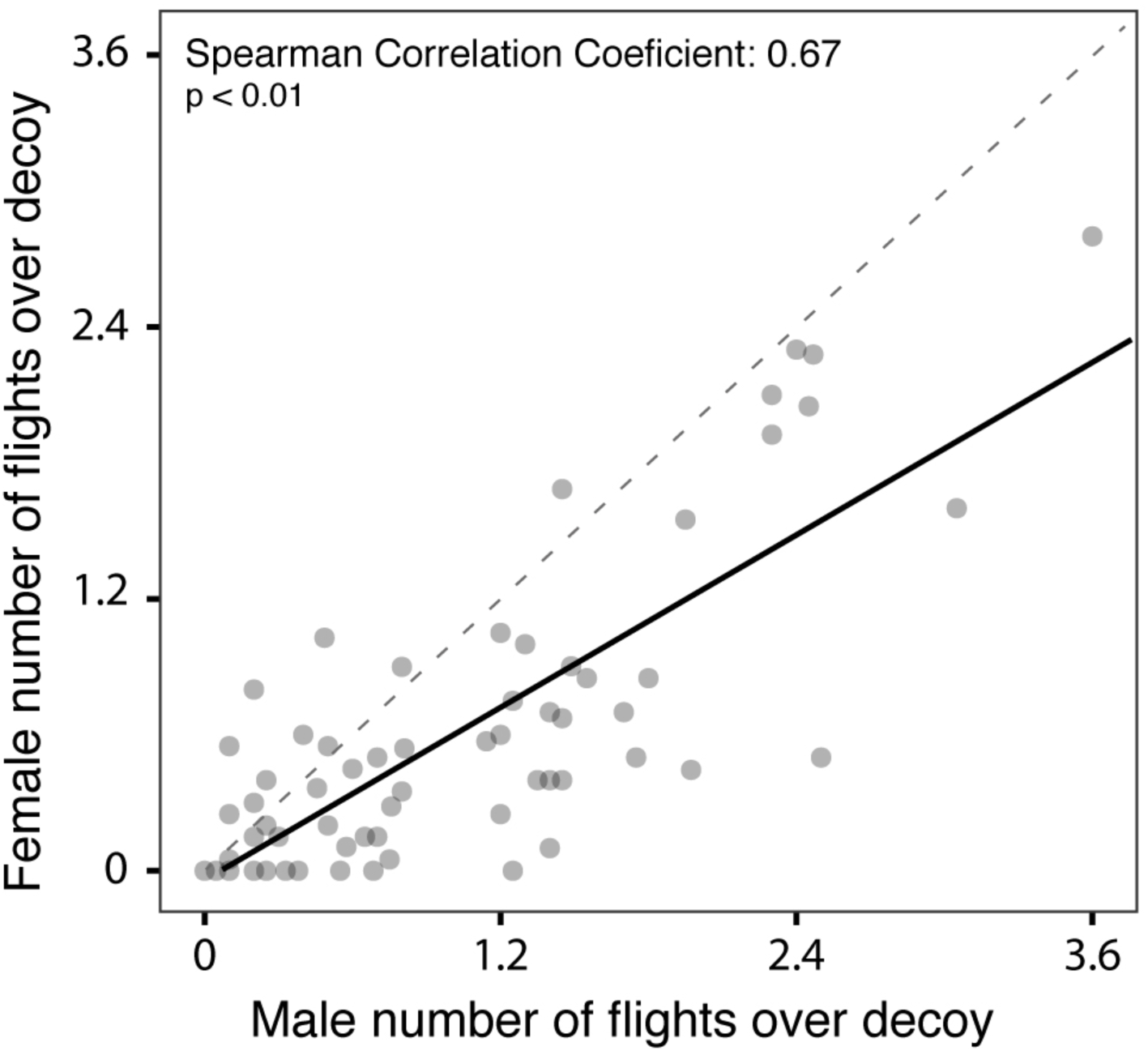
Female-male correlation of territorial defence, using number of flights over the decoy as proxy. The black line represents the regression line and the dashed line is the reference line with a slope of 1

## Discussion

We studied territorial defence in both male and female horneros and the role of duets during nest building and provisioning contexts. By using structural equation modelling, we were able to demonstrate that six observed behavioural variables (i.e. response latency, time spent within five metres of the decoy, average time spent on the nest during a visit, number of solo songs, number of duet songs, and number of flights over the decoy) were linked by an unmeasured latent trait “territorial defence”, both across sexes and contexts (Fig. 1). We also found that the number of flights over the decoy and duet songs were the variables with highest path loading to territorial defence. We then proceed to study independently those two traits with highest path loadings and showed that males were defending territories more strongly than were females during both breeding contexts, even though we only found weak evidence for differences between contexts within each sex (owing to large uncertainty in our estimate, Fig. 2). Lastly and as expected, we observed a strong positive correlation in territorial defence between males and females within the pair (Fig. 3).

The correlation structure of the latent variable territorial defence was similar in males and females, providing for the first time a sex-comparison of the latent variable “territorial defence” in birds. We also observed that the sign and magnitude of the path loadings between breeding contexts (i.e. nest building and provisioning) were very similar (Fig. 1B). Our results thus suggest the existence of a common structure in defensive behaviour during territorial intrusion in horneros, potentially underlined by a sex- and context-independent mechanism that is affecting all behavioural variables in a similar manner. However, to draw general evolutionary patterns of territorial defence it would be necessary to investigate whether the described latent variable is under selection as an integrated trait (i.e. functional module) and whether the same structure among traits is observed in other life-history stages (e.g. outside the breeding season) or in different ecological contexts (e.g. as territorial defence against predators or nest-parasitic species). Importantly, by using a structural equation modelling approach we were able to reveal complex relationships for multiple traits that would have otherwise not been possible to capture. Traditional statistical approaches such as multiple regression analysis or principal component analysis (PCA) are not suitable to evaluate different *a priori* defined hypotheses while accounting for trait correlation. PCAs are defined purely on the basis of mathematical associations between the traits and so their biological meaning can be challenging to interpret or even nonexistent. However, SEM provides a more interpretable method of viewing variation among correlated variables. Although SEM will ultimately be defined by mathematical associations like PCAs, with SEM, one makes use of biological information to fit the correlation structure to be tested among variables. Therefore, SEM has a clear advantage over PCA in terms of making biological inferences from the data. Lastly, another advantage of using a structural equation modelling approach is ralated to data collection methods. The fact that the number of flights over the decoy and duets songs were the variables with the highest path loadings to territorial defence indicates that by measuring only these two observable behaviours, and not all six, researchers should have a good estimation of territorial defence – at least in those studies that aim to quantify territorial defence towards conspecifics in horneros. Nonetheless, a good biological use of the approach would require a validation of the method for each new species where territorial defence is to be characterized.

In the context of territorial defence, duets have been mostly studied as a single trait independently of other complementary or related behaviours in multiple bird species (e.g. Hall and Peters 2008; Dowling and Webster 2016; Odom et al. 2017; Quirós-Guerrero et al. 2017). Here we explicitly tested for the first time whether duets are a behavioural variable linked by a latent trait, “territorial defence”. We did so by combining a classical STI approach with structural equation modelling. One of the predictions of the “joint territorial defence” hypothesis is that duets should play a more important role than solo songs (Hall 2009). As expected, in our study duets represented an important response during territorial defence for both sexes and breeding contexts. They were stronger than solo songs and were as relevant as other physical traits like the number of flights over the decoy. Our results are in line with previous findings in the species suggesting the territorial function of duets in hornero, overall providing evidence for the “joint territorial defence” hypothesis (Diniz et al. 2018, 2019, 2020). While previous studies tested this hypothesis by means of different approaches, the strength of our study resides in the application of a comprehensive method that accounts for the multivariate nature of territorial defence behaviours.

Males defended more their territories than females during both breeding contexts. At first glance, this is not surprising given that an unequal sex contribution of territorial defence has been previously reported in bird species that are socially monogamous and maintain territories year round (e.g. Willis 1972; Morton and Derrickson 1996; Bard et al. 2002; Fedy and Stutchbury 2005; Quinard and Cézilly 2012). In the specific case of horneros, however, male and female have been reported to contribute equally in most of the behaviours studied to date (Fraga 1980; Massoni et al. 2012; Diniz et al. 2020). However, there is strong evidence for sexual differences in singing-related traits independent of season (Diniz et al. 2018). Additionally, the observation that males engaged more in aggressive interactions than females has only been recently described (Diniz et al. 2018). Thus, our findings confirm and expand this observation with a standardized field test applied to a multivariate framework beyond song production. One explanation for the observed sex-differences might be a division of labour between members of a pair (e.g. Morton et al. 2000). For instance, males might invest more resources (i.e., time and energy) in actively defending their territory or nest (e.g. physical attacking the intruder), whereas females might focus on different activities (e.g. predator vigilance, guarding the nest against parasitic species). Another factor potentially explaining our results is that males and females might face different physiological (breeding) costs (e.g. Nilsson and Råberg 2001). In fact, during nest building (when females are close to egg laying) females have a poorer oxidative condition than males and are more sensitive to STIs, suggesting a sex-specific physiological cost of territorial defence (Mentesana and Adreani 2020). Lastly, our findings could also be influenced by the way the territorial intrusions were performed (i.e. with one single dummy). While plausible, this explanation seems unlikely given that horneros are monomorphic in body size and plumage colouration (Diniz et al. 2016) and the playbacks consisted of vocalizations from both sexes.

We did not find strong support for our prediction that the levels of territorial defence were higher during the nest building than in the provisioning context (Table 2). Given that extra-pair levels are very low in this species (∼3%, Diniz et al. 2019), one of the main assumptions of our prediction was that territory take-over was higher during the fertile period of the females, i.e., during nest building than during provisioning (Gill et al. 2007; Demko and Mennill 2018). It is possible that for horneros it is more beneficial to maintain constant levels of territorial defence in order to hold the territory year-round than the potential benefits of extra-pair paternity (Warner and Hoffman 1980). This might be especially the case when population densities are high, where comparable territorial defence can be expected across different life-stages as we observed in the horneros. Further research will help to shed light on these context-specific patterns.

Male and female aggression were strongly and positively correlated within the pair despite sex-specific differences in territorial defence. Our results are in line with previous findings of coordinated territorial defense on rufous horneros outside the breeding season (Diniz et al. 2020) and more generally with other studies showing that duetting birds were more collaborative within the pair than non-duetting species (see Logue 2005).Although our study cannot directly address the evolutionary relevance of pairs being positively correlated in their behaviours (e.g., fitness consequences), our findings suggest that exhibiting a joint territorial defence might be an important mechanism of pair bonding or pair stability (Wickler and Seibt 1980). In this direction, our study raises the question of whether pairs of horneros that show similar territorial defence levels would experience increased reproductive benefits (Schuett et al. 2010). Indeed, it is known from other bird species that pairs exhibiting comparably high levels of territory defence towards conspecifics attain higher reproductive success (e.g. in eastern blue birds, *Sialia sialis;* Harris and Siefferman 2014).Therefore, investigating patterns of selection on assortative mating in pairs of horneros poses an exciting avenue for future research.

## Conclusion

This work expands upon a classical body of research on territorial defence. We demonstrated that six observed behavioural variables quantified during a simulated territorial intrusion were linked by an unmeasured latent trait “territorial defence”. In particular, the number of flights over the decoy and the number of duet songs were the variables with highest path loadings to the latent variable “territorial defence”. Furthermore, this study fills an important gap in our knowledge about the role of duets. We provided support for the hypothesis that avian duets are a key component in the joint territory defence. Indeed, we showed that duets represented a stronger response of territory defence than solo songs, and that their importance was comparable to physical traits. Our study also highlights the importance of using more integrative, multivariate approaches to study behavioural traits. By applying a structural equation modelling framework, we were able to evaluate *a priori* hypotheses of how different behavioural variables were linked by an unmeasured latent trait. Such complex patterns would have not been possible to capture using traditional statistical approaches such as principal component analyses Hence, the combination of a classical behavioural approach like simulated territorial intrusions with structural equation modelling brings new exciting possibilities into the field of behavioural ecology.

## Authors’ contributions

NMA, LM and MM share first authorship and names are ordered at random. NMA and LM conceived the study and designed the study. BT provided logistic support. NMA, LM, EG and EC collected the data. NMA, LM, and MM analyzed the data. NMA, LM, and MM wrote the manuscript with input from all authors.

## Acknowledgments

We thank Klaus Pichler for his help to prepare the STI and all the “INIA Las Brujas” staff for supporting us with accommodation and equipment during fieldwork. We also thank Facultad de Ciencias and the Ethology lab from Universidad de la República and Juan Carlos Reboreda and the “Laboratorio de Ecología y Comportamiento Animal” at the Universidad de Buenos Aires for the logistical support, and Pablo Tubaro from the Museo Argentino de Ciencias Naturales ‘Bernardino Rivadavia’ (MACN) for providing us with the mounted hornero. We are grateful to Manfred Gahr and Michaela Hau for their valuable support. We also thank Yimen Araya-Ajoy, Glenn Cockburn, Luke Eberhart-Phillips and Wolfgang Wickler for constructive criticism on previous versions of the manuscript. Finally, we want to thank the anonymous reviewers for constructive feedback on the manuscript.

## Availability of data

The datasets generated and/or analysed during the current study are available in the open repository: Mendeley Data (https://data.mendeley.com/datasets/7ztwn539jd/1).

## Compliance with ethical standards

### Ethical approval

The experimental procedures of this study have approval by the Ethics Committee of Animal Experimentation (CEUA) of the Facultad de Ciencias of the Universidad de la República, Uruguay (Protocol number 186, file 2400-11000090-16).

### Funding

This work was funded by the International Max Planck Research School (IMPRS) for Organismal Biology, and by Idea Wild that provided field equipment.

