## Supplementary materials for "Defending as a unit: sex- and context-specific territorial defence in a duetting bird"

Journal: Behavioural Ecology and Sociobiology

**—Online Supplementary Material—**

**Table S1.** *A priori* hypotheses of behavioral integration between six measures of territorial defense.

| Model | Hypothesis |
| --- | --- |
| Model 1 | Null model of trait independence. |
| Model 2 | All six behavioral traits are linked by a latent variable “territorial defense”. |
| Model 3 | All behavioral traits are linked by a latent variable “territorial defense”, except duet during nest-building and duet during provisioning that are independent. |
| Model 4 | All traits measured during nest-building are linked by a latent variable “territorial defense during nest building”; and all traits measured during provisioning are linked by a latent variable “territorial defense during provisioning”. |
| Model 5 | All traits measured during nest-building are linked by a latent variable “territorial defense during nest building”; and all traits measured during provisioning are linked by a latent variable “territorial defense during provisioning”, except duet during nest-building and duet during provisioning that are independent. |
| Model 6 | All traits measured during nest-building are linked by a latent variable “territorial defense during nest building”; and all traits measured during provisioning are linked by a latent variable “territorial defense during provisioning”. There is an additional covariance between the two latent variables. |
| Model 7 | All traits measured during nest-building are linked by a latent variable “territorial defense during nest building”; and all traits measured during provisioning are linked by a latent variable “territorial defense during provisioning”, except duet during nest-building and duet during provisioning that are independent. There is an additional covariance between the two latent variables. |

**Table S2.** Results of model comparison using Akaike Information Criterion for small sample sizes (AICc) values to compare our seven candidate models. We constructed these seven models to test for “territorial defence” for each sex in both breeding contexts. To do so, the 12 different behavioural traits were modelled simultaneously. Smaller AICc values are given to models that better fit the data. Models whose AICc values differ from that of the top model (ΔAICc) by more than 2 are considered to lack explanatory power relative to the top model. The best supported model in both sexes was model 4, that hypothesized that all behavioural traits measured during nest-building are linked by a latent variable “territorial defense during nest building”; and all traits measured during provisioning are linked by a latent variable “territorial defense during provisioning”.

|  |  | **Model 1** | **Model 2** | **Model 3** | **Model 4** | **Model 5** | **Model 6** | **Model 7** |
| --- | --- | --- | --- | --- | --- | --- | --- | --- |
| **Male** | **AICc** | 184.9 | 161.8 | 175.7 | 147.7 | 157.3 | 150.5 | 159.8 |
|  | **ΔAICc** | 37.1 | 14.1 | 28.0 | 0.0 | 9.5 | 2.7 | 12.1 |
| **Female** | **AICc** | 226.2 | 185.9 | 197.1 | 139.4 | 177.9 | 142.4 | 180.1 |
|  | **ΔAICc** | 86.8 | 46.4 | 57.7 | 0.0 | 38.4 | 2.9 | 40.7 |

**Figure S1.** Models (1–7) of hypothesized relationships between six behavioral traits. Models are described in Table 1. Unidirectional arrows represent causal relationships between traits; bidirectional arrows represent undefined correlations. Solid lines represent relationships present across the whole set; dashed lines represent relationships expressed only in specific models. Dashed lines were active in model 2, 4 and 6; and absent (i.e., not modelled) in model 3, 5 and 7. Grey shaded boxes represent behavioural traits measured during provisioning contexts.

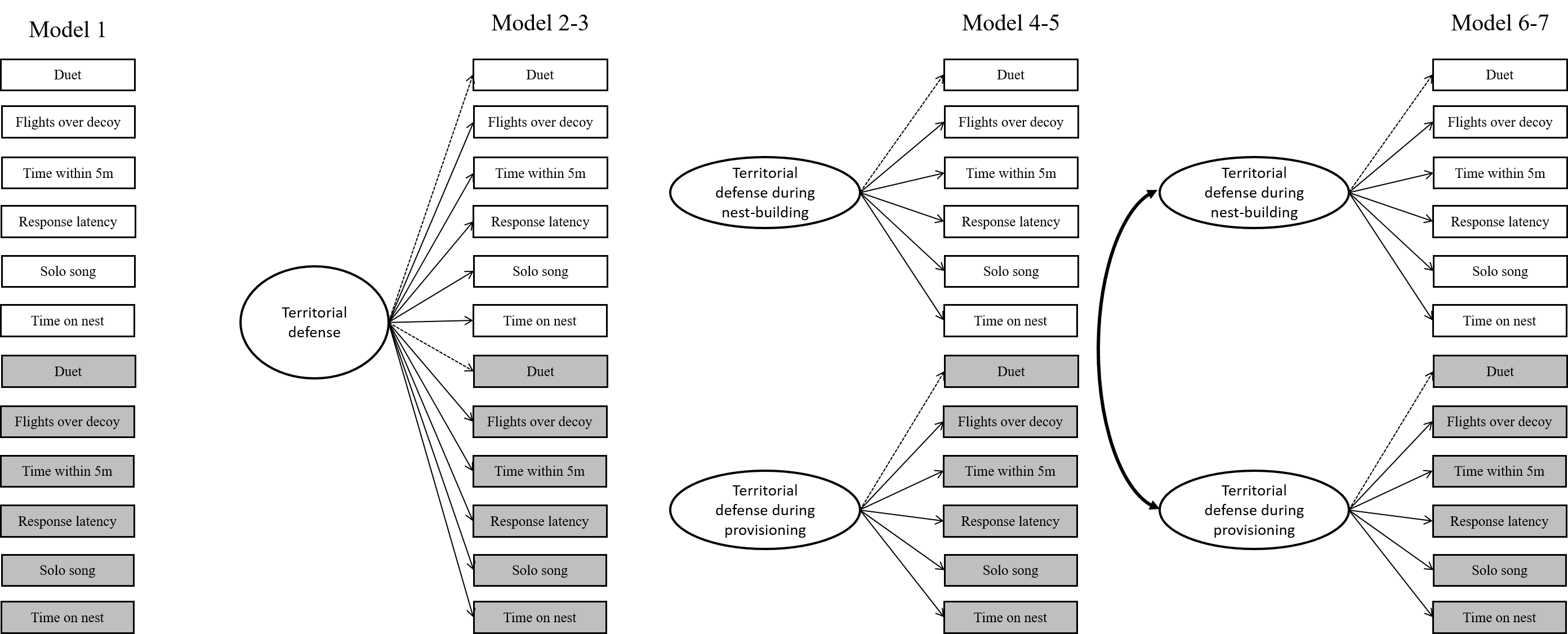

**Table S3.** Estimated phenotypic correlations for the six behavioural traits assayed during a simulated territorial intrusion in male horneros. We present among-trait correlations of individuals measured during the “nest building” context on the upper off-diagonals and during “provisioning”, on the lower off-diagonals. Sample size for each breeding context is shown in brackets.

|  | |  | Nest building (N= 39) | | | | | | |
| --- | --- | --- | --- | --- | --- | --- | --- | --- | --- |
|  | |  | | Time on the nest | Flights over decoy | Number of duet songs | Number of solo songs | Response Latency | Time within 5 m of decoy |
| Provisioning  (N = 25) | Time on the nest | | |  | 0.24 | 0.23 | 0.06 | 0.18 | -0.04 |
|  | Flights over decoy | | | 0.59 |  | 0.64 | -0.21 | 0.35 | 0.08 |
|  | Number of duet songs | | | 0.45 | 0.56 |  | -0.28 | 0.31 | 0.13 |
|  | Number of solo songs | | | 0.28 | 0.33 | -0.06 |  | 0.09 | 0.22 |
|  | Response Latency | | | 0.22 | 0.52 | 0.31 | 0.30 |  | -0.19 |
|  | Time within 5 m of decoy | | | 0.55 | 0.35 | 0.47 | 0.33 | 0.53 |  |

**Table S4.** Estimated phenotypic correlations for the six behavioural traits assayed during a simulated territorial intrusion in female horneros. We present among-trait correlations of individuals measured during the “nest building” context on the upper off-diagonals and during “provisioning”, on the lower off-diagonals. Sample size for each breeding context is shown in brackets.

|  |  | Nest building (N= 38) | | | | | |
| --- | --- | --- | --- | --- | --- | --- | --- |
|  |  | Time on the nest | Flights over decoy | Number of duet songs | Number of solo songs | Response Latency | Time within 5 m of decoy |
| Provisioning (N = 24) | Time on the nest |  | 0.24 | 0.16 | 0.40 | 0.03 | 0.07 |
|  | Flights over decoy | 0.22 |  | 0.79 | 0.20 | 0.42 | 0.47 |
|  | Number of duet songs | 0.06 | 0.65 |  | 0.27 | 0.48 | 0.42 |
|  | Number of solo songs | 0.25 | 0.39 | 0.31 |  | 0.25 | 0.25 |
|  | Response Latency | 0.04 | 0.35 | 0.35 | 0.02 |  | 0.16 |
|  | Time within 5 m of decoy | 0.17 | 0.51 | 0.77 | 0.46 | 0.19 |  |

**Table S5.** Parameter estimates (with standard errors, SE) from the best-supported model according to AICc model comparison.

|  | **Best-supported SEM model** | | | | | | | |
| --- | --- | --- | --- | --- | --- | --- | --- | --- |
|  | **female – nest building** | | **female - provisioning** | | **male - nest building** | | **male - provisioning** | |
|  | **Estimate** | **SE** | **Estimate** | **SE** | **Estimate** | **SE** | **Estimate** | **SE** |
| **number of solo song 🡨 Aggressiveness** | 0.31 | 0.17 | 0.48 | 0.21 | -0.02 | 0.19 | 0.41 | 0.21 |
| **number of duet song 🡨 Aggressiveness** | 0.75 | 0.15 | 0.87 | 0.18 | 0.72 | 0.20 | 0.64 | 0.20 |
| **flights over decoy 🡨 Aggressiveness** | 0.94 | 0.14 | 0.70 | 0.19 | 0.75 | 0.20 | 0.79 | 0.19 |
| **response latency 🡨 Aggressiveness** | 0.52 | 0.16 | 0.19 | 0.22 | 0.44 | 0.18 | 0.53 | 0.21 |
| **time spent on nest 🡨 Aggressiveness** | 0.21 | 0.17 | 0.12 | 0.22 | 0.28 | 0.19 | 0.73 | 0.19 |
| **time spent within 5m of decoy 🡨 Aggressiveness** | 0.58 | 0.16 | 0.92 | 0.17 | 0.03 | 0.19 | 0.64 | 0.20 |
| **number of solo song** | 0.90 | 0.21 | 0.77 | 0.23 | 1.00 | 0.23 | 0.83 | 0.25 |
| **number of duet song** | 0.43 | 0.13 | 0.24 | 0.13 | 0.48 | 0.23 | 0.58 | 0.20 |
| **flights over decoy** | 0.11 | 0.14 | 0.51 | 0.17 | 0.43 | 0.24 | 0.37 | 0.17 |
| **response latency** | 0.73 | 0.18 | 0.97 | 0.29 | 0.81 | 0.20 | 0.72 | 0.23 |
| **time spent on nest** | 0.96 | 0.22 | 0.99 | 0.29 | 0.92 | 0.22 | 0.47 | 0.18 |
| **time spent within 5m of decoy** | 0.67 | 0.17 | 0.16 | 0.13 | 1.00 | 0.23 | 0.59 | 0.20 |
